# Divorcing Strain Classification From Species Names

**DOI:** 10.1101/037325

**Authors:** David A. Baltrus

## Abstract

Confusion about strain classification and nomenclature permeates modern microbiology. Although taxonomists have traditionally acted as gatekeepers of order, the numbers of and speed at which new strains are identified has outpaced the opportunity for professional classification for many lineages. Furthermore, the growth of bioinformatics and database fueled investigations have placed metadata curation in the hands of researchers with little taxonomic experience. Here I describe practical challenges facing modern microbial taxonomy, provide an overview of complexities of classification for environmentally ubiquitous taxa like *Pseudomonas syringae*, and emphasize that classification and nomenclature need not be the one in the same. A move toward implementation of relational classification schemes based on inherent properties of whole genomes could provide sorely needed continuity in how strains are referenced across manuscripts and data sets.

## Confusion Abounds in Modern Bacterial Taxonomy

Communication between researchers is a foundation of all scientific disciplines, and clarity of the message is therefore essential for moving science forward. Alternatively, confusion of underlying messages leads directly to systemic problems and disagreements. For modern microbiologists perhaps the best example of how systemic confusion can slow research progress involves ongoing disagreements about bacterial classification and nomenclature, a confusion which is only amplified by the traditional entwinement of these two activities. The advent of ’big data’ has placed microbiology at a crossroads where we can either systematically change the way we think about describing strains or suffer within an ever expanding cloud of uncertainty. Now is the time to divorce classification of strains from any discussions about nomenclature based on species concepts and transition to a system based on genomic information, at least for metadata entry and to ensure continuity across manuscripts.

The root of this article lies in a frustration that many researchers deal with every day. A frustration born out of the clashes between how taxonomy should proceed in theory and the realities of how it proceeds in practice. Although nomenclatural confusion has always inconvenienced microbiology, the speed and focus of research as well as the dedication of large numbers of taxonomists previously enabled back and forth dialogues to smooth over ongoing disagreements. However, traditional taxonomic schemes have not efficiently dealt with the rapid influx of genomic data and were not designed to account for intrinsic challenges that arise when non-taxonomists publish metadata. Researchers have been arguing about bacterial species concepts since the dawn of microbiology [1-3], but the intent of this article is not to get caught up in discussions about what constitutes a bacterial species nor is it to suggest that the perfect mechanism for classification has been uncovered. My goal is to call attention to conflicts between the philosophy and practice of bacterial taxonomy, which generate much confusion for the classification of environmentally ubiquitious taxa like *Pseudomonas syringae.*

## The Philosophy of Bacterial Classification and Nomenclature

Taxonomy is a branch of microbiology that consists of three fundamental and often intertwined activities: identification, classification, and nomenclature of strains [4]. While these words can often be thought of as synonymous, important yet subtle distinctions can be drawn between these three areas. Whereas classification provides a means to index strains logically, it can exist independently of studies of how to accurately identify or name particular groups of strains. Therefore, taxonomy is not simply the study of organismal classification but a larger field that deals with the implementation of classification for diagnostic or naming purposes.

Bacterial strain and species names should relay useful information about properties or phenotypes generally shared by the groups that fall under defined nomenclatural umbrellas. One main reason for applying Linnean species names to microbes is that these names represent a shorthand for describing underlying aspects of the organism’s biology while also informing about classification. Farmers in the field or doctors in a hospital rely on accurate identification of strains in order to make informed decisions about how to proceed. In an academic context, although names are important for identification’s sake, they also provide continuity throughout manuscripts, and enable researchers to investigate their own systems and make general predictions from other work. As science and technology progress, and many researchers become beholden to large curated databases, metadata provides a way to rapidly access and apply information across taxa and studies. For instance, if metadata were perfectly applied, one need only search the words "*Pseudomonas syringae"* and all relevant references and datasets would be retrieved. This is currently not the case, as described below, because such a search would miss a variety of genomes and many publications that truly focus on *P. syringae.* The importance of proper curation of metadata to research is illustrated by the development of lines of investigation that employ text mining algorithms to discover emergent phenomena across systems [5]. How this metadata is curated significantly affects continuity and communication within manuscripts and across research groups.

In a perfect world, one would be able to see a name, garner something about the biology of that particular strain, and know exactly how other strains within the same assigned species name are related. For instance, *Helicobacter pylori* represents a cluster of strains which largely only lives within human stomachs and which are causative gastric ulcers and cancer [6-8]. Classification and nomenclature for *H. pylori* strains is straightforward, despite extensive diversity across strains, because the unique and specialized niche that this organism inhabits enables tight grouping of phenotypic and phylogenetic clusters [9]. In other cases, phenotypic traits of interest exist outside those normally used for classification so that additional modifiers are added. *Escherichia coli* is a common inhabitant of vertebrate gastrointestinal tracts, but certain modifiers are added to this species name to reflect important data about biology [such as uropathogenic *E. coli* (UPEC) enterohemorrhagic *E. coli* (EHEC), etc…] [10]. Naming schemes are by no means perfect and throughout the course of microbiology there have been situations where nomenclature and classification have pointed in opposite directions. These cases are useful because they highlight different requirements for microbial nomenclature between professional and academic settings, and they demonstrate how single classification schemes can function in one context but not the other. Even though all strains of *Shigella* cause similar disease symptoms in humans, there have been multiple independent evolutionary origins due to convergent evolution of phenotypes by different strains of *E. coli* [11]. This underlying evolutionary convergence might not matter much in a hospital, because, all else equal, doctors can prescribe similar treatments for all cases. However, conflation of different evolutionary lineages under the same name could affect how results are interpreted across comparative genomic studies unless researchers explicitly incorporate these phylogenetic nuances into their designs. While this particular example of confusion was clarified quite quickly, largely because of the importance of *Shigella* for human health, many other cases likely exist where nomenclature is wrong or misleading but which will languish in obscurity due to the lack of widespread interest.

## Nomenclature and Classification Schemes in Practice

When thinking about bacterial taxonomy, one cannot set aside historical momentum generated by the requirement of cultureability of strains in the early days of microbiology. The first step for any nomenclatural decision is traditionally the establishment of a "type" strain that is used to set a foothold for new species designations [12]. Following from cultureability, bacterial types are binned by observable properties at micro and macroscopic scales. Features such as staining with certain dyes (i.e. Gram stain), cell shape, flagellar type, nutrient profiles, and pathogenicity on certain hosts are still used to assign newly isolated bacterial strains into broad categories. One of the better known schemes today involves grouping of *E. coli* strains based on their O and H antigens (e.g. O157:H7)[13], while *Salmonella* strains have are typed according to their phage sensitivities [14]. Gradually, diagnostic assays have become increasingly specialized and technologically advanced, enabling finely grained classification of strains [15].

Although phenotyping schemes continue to work well in many cases, they can be time consuming and often require specialized equipment and expertise. Classification methods that use compositional properties of each genome, such as DNA-DNA hybridization, were developed to overcome these challenges [4]. Indeed, genomic hybridization remains a gold standard for species characterization. The creation of genotyping schemes was further welcomed because genomes were thought to reflect inherent properties of each strain and were potentially less prone to convergence or experimental variability. With the development of PCR, random genotyping methods such as Rep-PCR enabled closer views into the genetic relationships between or differentiation among strains and could be carried out at a lower cost and with less expertise required than phenotypic methods [4]. The DNA sequencing revolution sparked innovation of multiple layers of sequence based characterization, which ultimately focused on targeted sequencing and phylogenetic comparison of the 16s rRNA gene conserved throughout bacteria [16,17]. Paralleling the development of more specialized phenotyping methods, more finely grained comparisons of DNA sequences, such as multilocus sequence analysis (MLSA), promised increased resolution for classification within strains and species [18]. Finally, decreased costs and increased reliability of whole genome sequencing now provides an unprecedented amount of information for classification [3,4].

A crucial step for any formal taxonomic scheme involves some level of oversight, which today takes the form of dedicated groups of researchers that diligently curate and vet bacterial names [e.g. 19]. When a new bacterial species is proposed, vetting largely takes place by publishing a characterization of the type strain in the *International Journal of Systemic and Evolutionary Microbiology* [12]. Although there have been countless disagreements over what constitutes a bacterial species, it has been possible for these experts to take the time to reconcile the nomenclatural record. This system has held up quite well to scrutiny when phenotypes were of dire interest (as is the case of human pathogens), when there were numerous paid taxonomic positions whose purpose was to vet new entries into the nomenclatural arena, and when researchers could spend the time to learn ’a feel’ for their organisms. With funding cutbacks, and millions of new strains being identified yearly, it is unclear how sustainable this system will be going forward.

## Blindspots in Microbial Taxonomy

The history of the field has established a legacy where many firmly believe that phenotypic characterization should figure prominently in bacterial taxonomy [1,3,4]. This belief has culminated in what is referred to as the polyphasic approach to classification and nomenclature, where phenotypic characterizations are blended with genotypic and phylogenetic information to place strains into nomenclatural groups. This reliance on phenotypes for taxonomic purposes persists even though there are numerous cases where phenotyping is impossible for certain strains, such as those that cannot be cultured [17]. Instead of completely redesigning classification schemes to only incorporate molecular methods and sequencing, exceptions to nomenclatural rules have been created to account for outliers. For instance, when type strains cannot be cultured, a "Ca." is appended before the Linnean name representing the idea that these are only candidate taxa which have not been validated according to the established rules [12,20]. Furthermore, the ease of sequencing has democratized the taxonomic process, and anyone that can perform PCR can generate new data. New ’species’ are discovered by researchers who are not necessarily focused on taxonomy and have not been trained to classify these organisms. In these cases, when strains have not been fully vetted and assigned proper species names or where there is no type strain, the tradition is to simply place "sp." after the genus [12]. Lastly, with the increasing number of strains that are sequenced and cultured, the more exceptions that arise to the established phenotypic rules for any particular group. What may have traditionally fit into a nice taxonomic box based on phenotyping methods may falter now, even for well described taxa, because exceptions or complications frequently arise with greater sampling [21]. Furthermore, with greater scrutiny, some already established phenotyping schemes fall apart due to inherent variability of the phenotypes themselves within single strains [22].

Taxonomists continue to struggle with how to incorporate phenotypic and genotypic methods for bacterial classification and to how to name bacterial species [1,3,4,23]. This reliance on human subjectivity builds in an inherent weakness for continuity across data sets and manuscripts. Every time a new phenotype of interest has emerged, or every time a lineage is split due to actual differences in evolutionary signals, there is a possibility that names of species or even genera change. While this practice is no doubt important from a taxonomic standpoint, the plasticity of nomenclature presents a significant obstacle for chasing down relevant data in historical literature because information about strains of interest is often hidden by changes in name for each strain. Advances in publishing technologies also work against the traditional nomenclature schemes. Although every new species name should be thoroughly documented and vetted with publication in designated journals, with the proliferation of modern journals it is easier than ever to publish papers regardless of their conclusions. Reviewers’ time is limited, and many may be unaware of current nomenclatural challenges, so that papers may be published where strain descriptions conflict with previously agreed upon taxonomic standards. Such situations amplify confusion in the literature because lineages are then officially referred to in multiple ways without a clear way to delineate the official taxonomic status.

## The *P. syringae sensu lato* Species Complex as an Example

There is no better system to illustrate the nomenclatural challenges of modern day microbiology, and to give examples for confusion arising from the points mentioned above, than *P. syringae.* Pseudomonads can be found ubiquitously across environments, but are also well known as pathogens of humans, animals, insects, fungi, and plants [24]. Therein lies one of the greatest challenges of pseudomonad taxonomy, that nomenclature within this group has often been biased by relying on phenotypes related to pathogenicity across hosts. It is possible, and even likely, that pathogenic contexts represent only a small sample of the selective pressures faced by strains in nature [25]. Therefore, phenotypes that might be of interest in defining species names may not be those that play a large role in the lifestyle of the strains and might therefore not lead to the most cohesive groupings. In a way, *Pseudomonas* is the taxonomic opposite of *H. pylori.*

The name *Pseudomonas syringae* began as a formal way to classify pseudomonad pathogens of plants, with the name "syringae" coined because the original type strain was a pathogen of lilacs within the genus *Syringa* [23]. Eventually, the LOPAT diagnostic scheme was created to classify strains as *P. syringae* based upon five different phenotypes; levan production, oxidase, potato soft rot, arginine dihydrolase, tobacco hypersensitivity. Species level nomenclature was layered onto this scheme, usually defined by pathogenic reactions and symptoms on host plants. Given the intrinsically changeable nature of species names, strains broadly classified within *P. syringae* are also currently informally grouped into the designations *sensu lato* (the largest sense) and *sensu stricto* (the smallest sense). *P. syringae sensu lato* includes all species that could be classified as part of *P. syringae* according to the LOPAT scheme genomic comparisons but in the absence of any other phytopathogenic information. *P. syringae sensu stricto* includes only those strains that are closely related to and phytopathogenically similar to the original type strain. After bacterial species nomenclature was overhauled in 1980, this system was modified to include "pathovar" or "pathogenic variety" status as a way to reflect previously established boundaries between strains with distinct phytopathogenic profiles [26]. Although the causes are outside the bounds of this piece, strains within *P. syringae sensu lato* have been split up into multiple species with subsets assigned pathovar designations. Some pathovars themselves are polyphyletic due to convergence in host range (i.e. pv. *avenellae)*, whereas others are tightly grouped (i.e. pv. *phaseolicola)* [26,27]. There are even some lineages known to cause disease and which are model systems for host pathogen interactions, but which are not part of a monophyletic species cluster (i.e. *P. syringae* pv. *tomato* DC3000)[23]. As with many other bacteria, taxonomy of *P. syringae* has increasingly relied upon on genomic properties and DNA sequences. Strains within the group *P. syringae* were first categorized and split up into nine different genomospecies based upon DNA-DNA hybridization [24]. Later, strains were classified into 13 different groups by multilocus sequence type ( MLST) and multiple subgroups, with a reduced yet accurate scheme including the loci *rpoD, gltA, cit*, and *gyrB* [28,29], Whole genome phylogenies can now be built using a variety of methods and with draft genomes, all approximate the earlier classification schemes (although there are notable exceptions) [30-32]. Given the current state of the field, one can reasonably determine where within *P, syringae sensu lato* their strains of interest fall given a sequence for only the *gltA* locus (Figure 1 and [28]).

Perhaps the greatest challenge with taxonomic classification of *P, syringae* is the broad phenotypic versatility and variability that strains display. This bacterium is an environmentally ubiquitous hemibiotroph that can be isolated from environmental sources including rivers, lakes, leaf litter, and clouds [33-35]. For any given strain, host range can be difficult to catagorize because the occurrence and severity of symptoms is dependent on the precise environmental conditions and sensitivity to these conditions is itself influenced by phylogenetic placement [36]. Moreover, just because strains fail to cause disease does not mean that they will fail to grow *in planta* or survive epiphytically on those hosts. Adding to the confusion, numerous strains have been isolated from environmental resources and from asymptomatic plants [21]. Many of these environmental strains are closely related to known pathogens of plants, and may even be capable of causing disease under laboratory conditions, but these are not given pathovar designations because they are non-pathogenic or their host range has not been determined [39,40]. To these points, the main reason that genomospecies have not been granted official taxonomic status is because there are no consistent phenotypic differences that differentiate them.

Since *P, syringae sensu lato* has become a model system for environmental genomics, the genome sequences of hundreds of individual strains have been deposited in publicly accessible databases [41,42]. The inconsistency of metadata across these strains makes it difficult, in the least, to mine much of the information that would be useful for comparative genomic analyses. Numerous cases currently exist within Genbank of strains that could be classified in the *P, syringae* species complex that are arguably mischaracterized as an incorrect or obsolete species name. Therefore, if one is interested in accessing genomes for all of these strains, they either have to search and find which species name the genome is listed under or perform repetitive sequence based searches to gauge how strains are related. These inconsistencies hamper research because they prevent researchers from discussing overlapping data sets and lead to wasteful redundancy when multiple groups are unaware that they are independently investigating the same strains.

This already confusing taxonomic situation has been compounded by a variety of published papers which refer to strains with old names, rename strains incorrectly, or differentially name the same strain because of grey areas in nomenclature. One of the best examples involves a strain from pathovar *phaselicola*, which is a pathogen of green beans [26]. This strain has variably been referred to as *P, syringae* pv. *phaseolicola* or *Pseudomonas savastanoi* pv. *phaseolicola* even though it could properly be named as *Pseudomonas amygdali* pv. *phaseolicola* according to current rules [23,26,43]. There are also numerous cases where strains currently exist in taxonomic limbo as *Pseudomonas* sp. because they have not been formally described (i.e. UB246 in Figure 1) [21,28]. The widespread challenges of properly referring to strains within this species complex are demonstrated in Figure 1. Even though most of the strains represented in the figure can be referred to as *P. syringae*, as shown by the highlighted names, multiple other species names are currently used by different research groups. Despite the best intentions of all researchers, nomenclature remains inconsistently applied across these strains. It would be one thing if the taxonomic confusion were limited to one particular strain within *P. syringae sensu lato*, but such confusion is seemingly universal and widespread across lineages. I have no doubts that the situation is equally as confusing across other environmentally ubiquitous groups that contain phytopathogens (i.e. *Burkholderia, Erwinia, Rhizobium* [44-46]), or across groups like *Vibrio* where conflicts arise between traditional nomenclatural schemes and in depth analyses of population genetics signals [47].

## Divorcing Strain Classification from Species Names

The ever-increasing flood of genomic data will lead to an increase in nomenclatural confusion across taxa. New DNA sequencing technologies are continuing to emerge and mature so that, very soon, direct sequencing of nucleotides and single cell genomics may be possible under field conditions [47]. Complete genome sequencing will eventually be cost efficient and straightforward enough to use for rapid classification across all taxa, even the uncultureable majority. Along these lines, it is worth noting that genome sequencing has potentially overtaken phage phenotyping as the preferred method for *Salmonella* classification in the United Kingdom [48]. Although phenotyping will certainly still be useful for more thorough studies, these tests will always be likely used in combination with genotypic data. Furthermore, it is inevitable that researchers who are only casually interested in proper taxonomy will continue to submit sequences to databases and to publish papers. In this light, what can be done to minimize future confusion?

A key to clearing these challenges is the realization that taxonomic classification and nomenclature are not commutative. Although the ability to name strains inherently requires classification, classification itself can take place sequentially based on inherent genomic properties and through comparison to strains that are already indexed without the need to group strains together under potentially subjective nomenclatural umbrellas. All that is necessary for such a system is agreement as to an algorithm for classification, and the ability to identify the most closely related strain according to this scheme. Furthermore, although discussions about how to name bacterial strains inevitably deteriorate into disagreements about how to define bacterial "species", establishment of an independent classification scheme would separate these discussions and disagreements from indexing these strains in practice for posterity.

There will never be a perfect classification system, but we can develop a scheme centered around whole genome sequences which ensures continuity of research messages across databases and papers. To achieve any level of success this scheme must be relational in that new designations are assigned solely through genotypic similarity to previously indexed strains whether or not they have been formally characterized as a type strain, and it must exist independently of disagreements about species concepts. An added benefit of relational classification is that identifiers could also provide information about relationships between strains. This system must be expandable so that discoveries of new clades do not force a complete rewriting of previous classifications, and must be applicable to uncultured strains. Classification by this system must be automatable and function well regardless of level of taxonomic expertise, and could even be retroactively applied across all published genome sequences and manuscripts. Ideally, these classifications would already be implementable given the infrastructure of Genbank and could be added to keywords of journal articles to streamline search strings for text mining software.

Numerous options for such a classification scheme already exist, they just need to be universally implemented. MLSA/MLST or 16s rRNA schemes work well except they can be confounded by recombination, can be biased in the subsets of loci used, and ultimately just contain a lower number of informative sites than whole genomes [3,4,17,18]. Moreover, assigning group numbers based on static numbers significantly decreases the ability to efficiently add intermediate taxa at later times and limits the capability to relay information about relatedness between strains through relational classification. For instance, MLST group 2 in *P. syringae* has already been expanded to include MLST groups 2a/2b/2c/2d but what would happen if group 2a were further split? Likewise, *P. syringae* MLST group 10 is more closely related to group 5 than to group 9 according to the current typing scheme [28]. Life Identification Numbers (LINS) have been suggested as an alternative scheme that fits this bill, whereby the average nucleotide identity (ANI) of genomes of interest are sequentially compared [49,50]. After one genome is designated as a reference point (denoted by inclusion of 0s at all positions in its identifier), new genomes are compared and assigned standard numerical classifications based on overall genome similarity to the most closely related genome already assigned a LIN. Each position within the LINS identifier (denoted by subscript letters as in Figure 1) represents an increasing threshold of similarity for ANI between two strains. According to this scheme, strains that differ in number at position "A" have a much greater difference in ANI than strains that differ in number at position "E". Assigning identifiers to new genomes is akin to a first order Markov process, because classification is memoryless and based solely on the most similar genome already classified. In order to assign LIN values to a strain of interest one only needs to identify the most closely related previously categorized strain and compare ANI values. Due to the sequential nature of this scheme, increasing thresholds can be utilized where necessary in order to delineate closely related taxa, which would be manifest as additional positions added to the identifier (i.e. 0A,0B,0C,0D,0E,1F/0A,0B,0C,0D,0E,0F instead of 0A,0B,0C,0D/0A,0B,0C,0D). Firm borders between species groups need not exist because schemes like LINS provide progressive classification where one simply includes more finely grained comparisons of genome similarity. LINS is a computational example of the goal of moving to classification based on DNA hybridization, except that classification can be automated and take a fraction of the time and expertise. Notably, this system has been applied to strains within *P. syringae* [51].

Some have highlighted the weaknesses of classification solely through genome sequences [1,3,4,50]. Substantive challenges include instances where whole genome data does not exist, but strains can be partially assigned based on a limited set of genomic characteristics (such as MLST or 16s rRNA loci) where necessary. There will also be challenges for strains which have experienced extensive genomic diversification, either through selection or drift, as is the case with genome minimization across many intracellular parasites [51]. Bull *et al.* also bring up the idea that strains are often mislabeled or mixed up during storage or sequencing [4]. However, such problems exist regardless of classification scheme and an automated system of classifications linked to publication would actually enable interested parties to track down incorrectly named strains. Lastly, one of the greatest weaknesses to new classification schemes is not the most rational objection, but is nonetheless important to consider. Many researchers have become comfortable with and have grown accustomed to assigning binomial names to new microbes. Numerical codes are a less elegant way to describe new taxa and will be inherently unpalatable to some, regardless of whether this is implemented in parallel with the binomial system.

## Concluding Remarks

The overall message of this piece is not to throw out all previous taxonomic systems and start anew, but that we must move to implement a sequence-based classification system that is unambiguous. We can create a retroactive and expandable system that could be used by regulatory agencies, in publication keywords, and with metadata that exists independently of species nomenclature or concepts. This system would enable microbial classification to remain unchanged going forward. These systems can be expandable, with algorithms that can be created to automatically classify or group genomes by similarity when they are submitted to databases or to journals. This suggestion parallels both the call for universal DNA Barcoding across species [52] as well as the creation of ORCID identifiers (orcid.org) to trace publications by individual authors. Researchers from across disciplines will likely admit that something has to be done to change the way strains are classified, challenges which are exemplified quite well by *P. syringae sensu lato* (Figure 1). There may not be one single answer to the problem of nomenclatural confusion in microbes, but the today’s challenges will only worsen unless we rethink taxonomic strategies.

**Figure 1.**
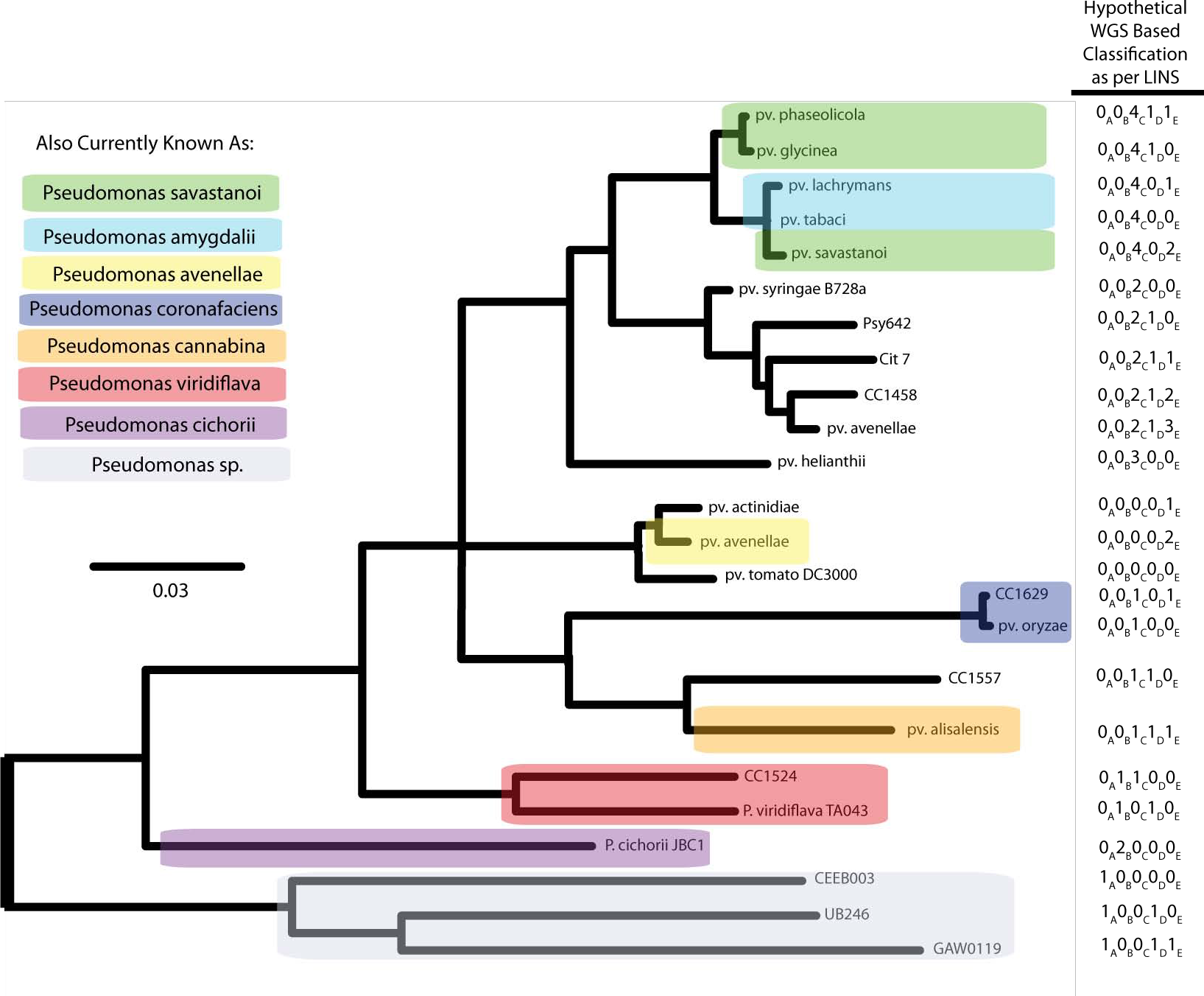
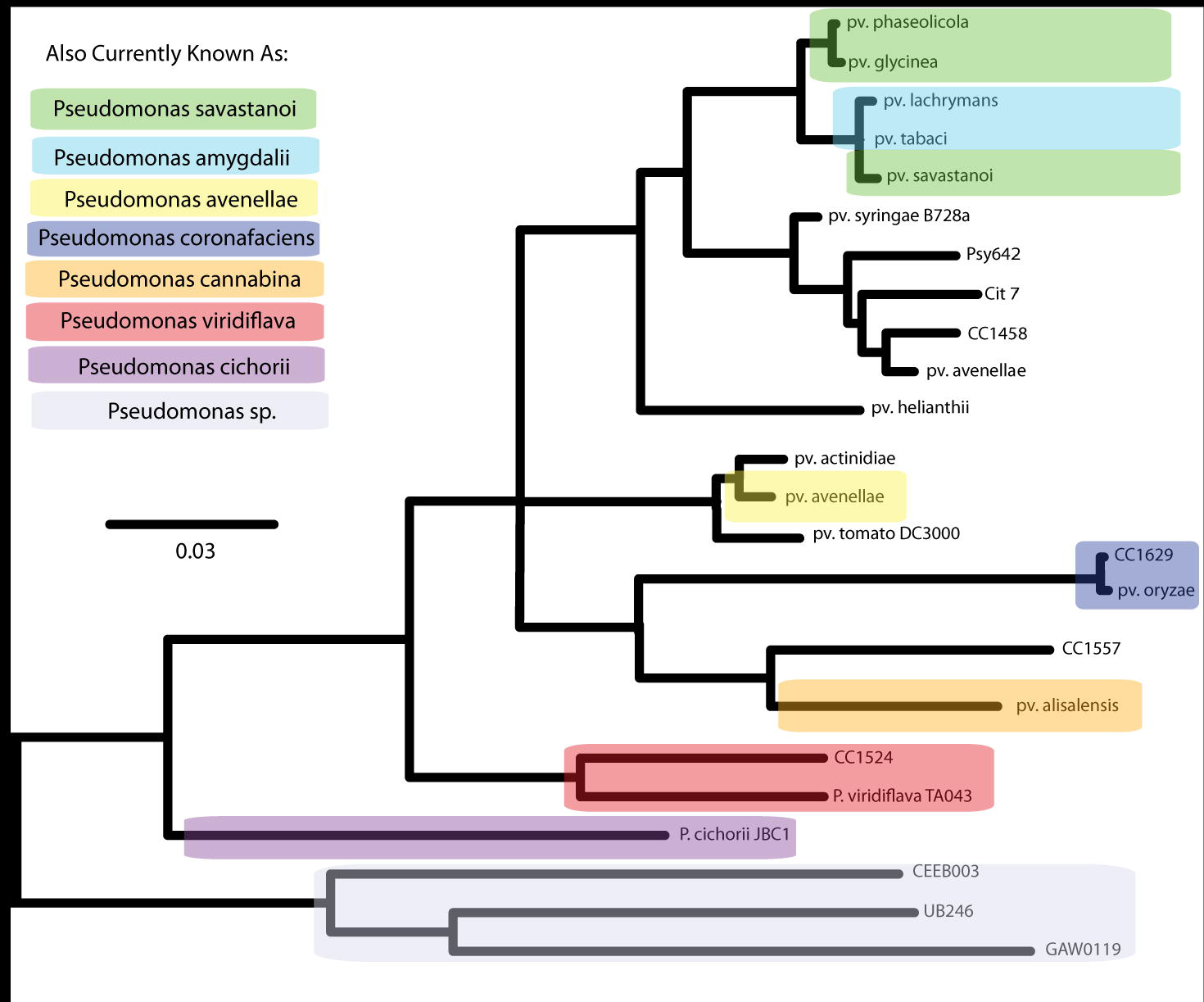
Current and Potential Classification of *Pseudomonas syringae sensu lato*. A Bayesian phylogeny was built using the full-length sequence of *gltA* from a diverse subset of *P. syringae sensu lato* genomes. Although each of these strains can be referred to as *P. syringae* in published papers and in metadata, strains that are currently referred to by species names other than *P. syringae* are highlighted in different colors. Strains which are known pathogens are labelled as either "pv." for pathovar or with alternative species names in the absence of pathovars. Strains which have been isolated from environmental sources or from plants in the absence of official disease classification have been labeled only with the strain number. To the right of each strain name is a mock numerical classification, based upon an example scheme described in [49], but in the absence of defined ANI thresholds or actual genome comparisons. Each column within this identifier represents an increasing threshold for ANI between that particular strain and the most similar genome already assigned a LIN. For instance column A could represent an ANI of 40%, followed by column B at 50%, C at 75%, D at 90%, E at 99%. Different numbers within these columns represents that particular strains differ from a common reference for that threshold ANI. Abbreviations: WGS, whole genome sequencing; LINS, Life Identification Numbers; ANI, Average nucleotide identity.

